# IDOL deficiency inhibits cholesterol-rich diet-induced atherosclerosis in rabbits

**DOI:** 10.1101/2022.05.03.490218

**Authors:** Tomonari Koike, Yui Koike, Yanhong guo, Dongshan Yang, Jun Song, Jie Xu, Xiangjie Zhao, Tianqing Zhu, Ruiting Li, Bo Wen, Duxin Sun, Oren Rom, Renzhi Han, Jianglin Fan, Minerva T. Garcia-Barrio, Jifeng Zhang, Y. Eugene Chen

## Abstract

**BACKGROUND:** The E3 ubiquitin ligase IDOL (Inducible Degrader of the LDL-Receptor) contributes to regulation of cholesterol metabolism through degradation of LDLR, VLDLR and ApoER2. Human genetic studies support the hypothesis that IDOL could serve as a target for the treatment of dyslipidemia. However, species-specific differences in overall lipid metabolism and IDOL regulation require new preclinical models to realize its therapeutic potential. We leveraged the advantages afforded by the rabbit model to address those limitations and generated a novel rabbit IDOL knockout, which we characterized in the context of atherosclerosis.

**METHODS:** *IDOL*^-/-^ rabbits were generated by CRISPR/*Cas9* technology. *IDOL*^-/-^ and wildtype littermates, on standard (SD) and atherogenic high-cholesterol diets (HCDs) were compared through assessment of lipid and lipoprotein profiles, triglyceride clearance, lipoprotein lipase (LPL) activity, liver pathology, atherosclerosis development, and fecal cholesterol, with bile acid contents assessed by mass spectrometry.

**Results:** Hepatic IDOL expression was increased in response to hypercholesterolemia and hypertriglyceridemia induced by HCD. On SD, loss of IDOL increased LDLR stability with reduced total cholesterol in plasma. On HCD, *IDOL*^-/-^ rabbits showed simultaneous and remarkable reduction in hypercholesterolemia and hypertriglyceridemia associated with enhanced lipid clearance and LPL activity as well as increased bile acid excretion in feces. *IDOL*^-/-^ rabbits presented markedly reduced HCD-induced atherosclerosis in the aorta and left coronary artery, without enhanced liver steatosis.

**CONCLUSIONS:** Loss of IDOL in rabbits recapitulates human genetic findings, thus setting the stage to accelerate preclinical studies towards development of strategies targeting IDOL for the treatment of atherosclerotic cardiovascular disease.

## INTRODUCTION

Recent therapeutic advances considerably reduced the incidence of cardiovascular disease (CVD) in the context of hyperlipidemia. The contribution of low-density (LDL) and very low density (VLDL)-lipoproteins is critical in atherogenesis.^1,2^ Clinical trials and meta-analyses established that LDL-C lowering, particularly through statins, reduces the progression of coronary atherosclerosis and the risk of coronary events.^3,4^ Mechanistic findings, clinical genetic epidemiology and drug targeting, led to combination therapies involving a statin and non-statin agents that are highly effective in patients with moderate-to-high cardiovascular risk.^5-10^ Regardless, for a significant proportion of statin-treated patients, with or without combination therapy, insufficient LDL-C reduction and relatively high residual risk from persistent high triglyceride (TG) levels still remain,^11,12^ thus limiting the benefits of these therapies^13-24^ and underscoring the need for new therapies targeting lipid metabolism for CVD prevention and treatment. Initial approaches using LXR agonists ameliorated atherosclerosis and a wide range of inflammatory disorders in preclinical animal models, yet failed Phase I clinical trials so far due to side effects, including hepatic steatosis and hypertriglyceridemia.^25,26^ Species-specific differences in liver LXR-dependent upregulation of Inducible Degrader of the LDL-Receptor, *IDOL* (a.k.a. *MYLIP*)^27,28^ could also contribute to those adverse outcomes through degradation of hepatic LDLR and ensuing LDL-C increase in plasma, as demonstrated in non-human primates.^29^

IDOL is a widely expressed E3 ubiquitin ligase induced transcriptionally by LXR/RXR in response to increased sterol signaling.^28,30^ Three IDOL targets, LDLR, VLDLR and ApoER2, are critical regulators of cholesterol and triglycerides metabolism.^31^ IDOL-mediated ubiquitination of the cytoplasmic tail of those receptors promotes their lysosomal degradation.^32-34^ Over a decade of research supports the potential of IDOL-based therapies for regulation of dyslipidemia.^35-37^Lack or down-regulation of IDOL increased LDLR stability, independently of SREBP or PCSK9.^38,39^ Conversely, liver-specific expression of dominant-active IDOL increased hypercholesterolemia and exacerbated atherosclerosis in mice.^40^ Human genetic studies found that reduced expression or loss-of-function polymorphisms in IDOL were associated with lower total cholesterol (TC) and LDL-C, and reduced CVD, supporting the feasibility of IDOL-based therapies.^41-46^ Unfortunately, critical differences in IDOL regulation in the liver between mice and primates (humans and monkeys) interfere with expedited clinical translation of the findings.^29^ Rabbits better approximate human lipid metabolism overall, providing better models in preclinical settings for CVD research.^47,48^ Whereas *Idol* is not an LXR-target in the mouse liver,^29,49^ here we show that *IDOL* expression is increased in rabbit liver upon LXR-agonist induction, as in non-human primates.^*29*^ HCD also increases hepatic *IDOL* expression, suggesting a direct role of IDOL in liver-mediated plasma lipid clearance. We demonstrate that knockout of *IDOL* (IDOL KO) increased the abundance of its targets and, importantly, reduced hyperlipidemia without enhancing liver steatosis. Consequently, loss of *IDOL* protected rabbits against HCD-induced atherosclerosis. This work contributes novel insights into IDOL biology and provides a new animal model relevant to human physiology to reinvigorate and expedite preclinical efforts towards IDOL inhibition as a potential therapeutic target for atherosclerosis-associated CVD.

## METHODS

The data that support the findings of this study are available from Dr. Y.E. Chen upon reasonable request.

### Generation of IDOL KO rabbits, husbandry and atherosclerosis model

New Zealand White rabbits (Covance Inc. Princeton, NJ) were used for this study. The *IDOL* knockout (IDOL KO, for the model; *IDOL*^-/-^, for the genotype) rabbits were generated by CRISPR/Cas9 as described previously,^50^ using a guide RNA (gRNA, GGATCTCCCAGCAGATGGACggg) targeting exon 2 (Figure S1A-B). Rabbits were housed individually in cages under constant 20°C temperature and 12-hour light/dark cycles. Rabbits were fed 120g/day of either a standard diet (SD, Teklad global rabbit diet 2030, Envigo, Indianapolis, IN) or a high-cholesterol diet (HCD, custom diet containing 0.3% cholesterol and 3% soybean oil added to the SD, Envigo) for 16 weeks to induce hyperlipidemia and atherosclerosis.^51^ At endpoint, the rabbits were fasted overnight, anesthetized with 4%–5% isoflurane inhalation and euthanized by exsanguination and thoracotomy.^52^ All animal studies were performed in accordance with the animal protocols approved by the Institutional Animal Care and Use Committee (IACUC) at the University of Michigan. Further details in online Supplemental materials.

### Statistical analysis

All data are expressed as mean±SEM. Statistical analyses were performed using GraphPad Prism 9, GraphPad Software, San Diego, CA. Statistical analyses for comparison between 2 groups or among 3 groups were selected based on the Shapiro-Wilk normality test for the distribution of each data set. For 2-group comparisons, two-tailed, unpaired Student’s t-test (parametric) or Mann-Whitney U-test (nonparametric) were used. For ≥3-group comparisons, we performed ordinary one-way ANOVA and Tukey’s multiple comparisons test (parametric) or Kruskal-Wallis and Dunn’s multiple comparison test (nonparametric). Two-way ANOVA and Sidak’s multiple comparison test were performed for data obtained from multiple experimental time points or conditions. Repeated measures (RM) Two-way ANOVA was used for analysis of repeated biweekly measures of plasma lipid changes. Statistical significance was set at P < 0.05.

**Other methods** can be found in the online Supplemental materials.

## RESULTS

### The LXR/IDOL/LDLR axis is active in the rabbit liver

To determine the suitability of the rabbit model to study the role of IDOL in lipid metabolism and atherosclerosis, we first tested whether the rabbit liver recapitulates the primate response to LXR agonists using the synthetic LXR pan agonist T0901317.^29, 53^ The agonist (10 mg/kg/day, 4 days) increased *IDOL* mRNA expression in the liver by 2-fold (Figure 1A), comparable to data from the cynomolgus monkey studies.^29^ Although the IDOL protein cannot be satisfactorily detected with the current commercial antibodies, we determined that the increase in *IDOL* mRNA resulted in 43% reduction in LDLR protein abundance, the primary target of IDOL in the liver, indicative of increased IDOL function (Figure 1B). To provide further validation of the rabbit model, we fed rabbits an atherogenic HCD for 6 weeks. At endpoint, *IDOL* expression in the liver of rabbits on HCD was increased by 2.8-fold, compared with the rabbits fed a SD (Figure 1C). These findings are in contrast with the limited sterol-dependent regulation of *Idol* expression in the mouse liver,^29^ indicating that the LXR/IDOL/LDLR axis in the rabbit liver resembles that of primates.

**Figure 1.**
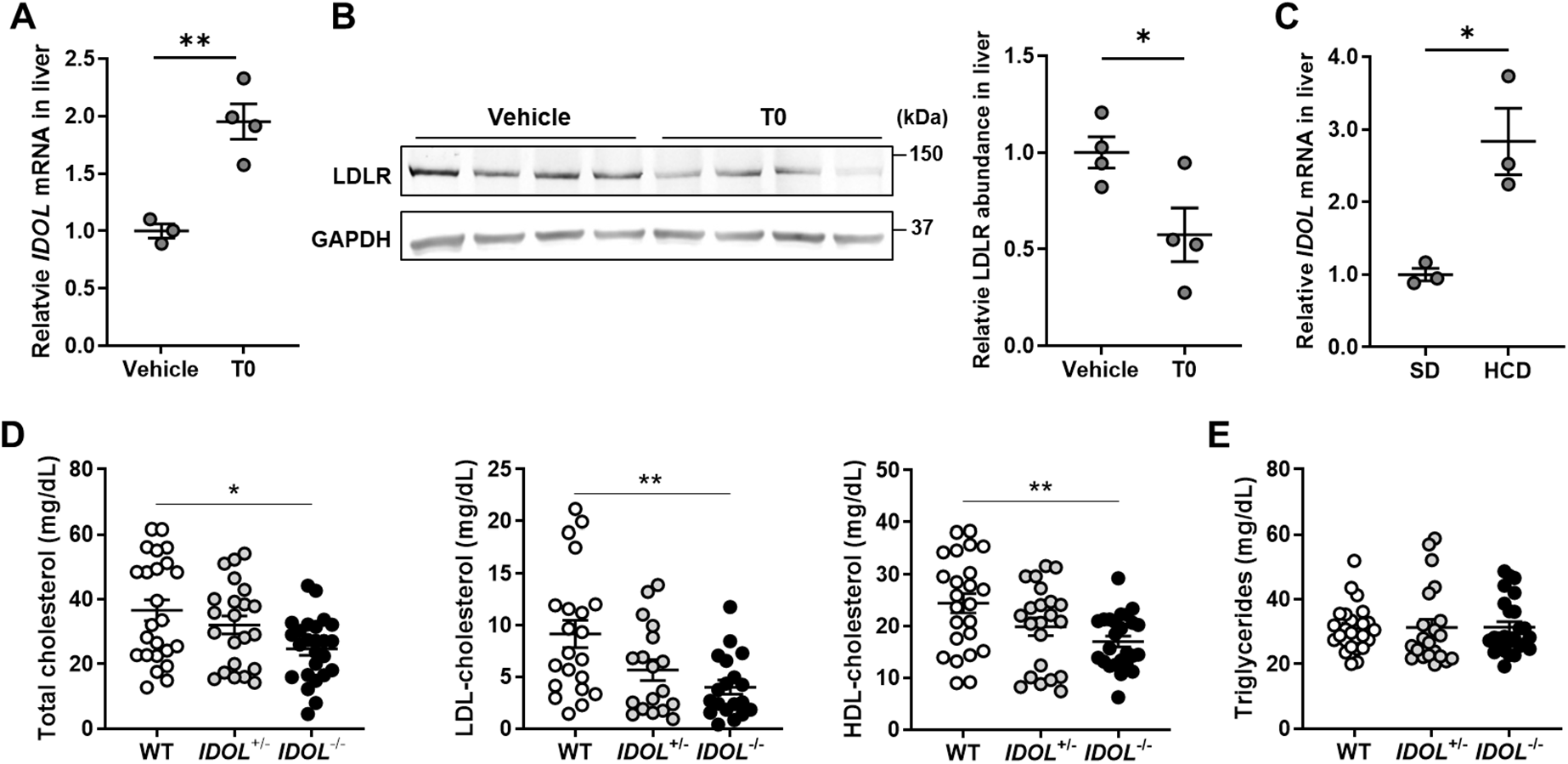
The LXR/IDOL/LDL axis is active in rabbit liver and IDOL KO reduces cholesterol in standard diet. **A**, New Zealand White (NZW) rabbits (5-months old, male) were administered the pan LXR agonist T0901317 (T0, 10 mg/kg/day) (or equivalent amount of DMSO, vehicle control) for 4 days and euthanized 4h after the last i.p. injection (n=4). Gene expression of the Inducible Degrader of the LDL-Receptor, *IDOL*, in the rabbit liver by qPCR and normalized by 18S RNA. Vehicle was set as 1. **B**, Western blot for Low-Density Lipoprotein Receptor, LDLR, in the liver of rabbits as in A (left), normalized by Glyceraldehyde-3-phosphate Dehydrogenase, GAPDH. Densitometry analysis (right) was performed using Image J (n=4). Vehicle was set as 1. **C**, *IDOL* mRNA expression, determined as in A, from livers of rabbits on a standard diet (SD) or a high-cholesterol diet (HCD) for 6 weeks (5-months old, male, n=3). SD was set as 1. **D**, Total cholesterol (TC); LDL-cholesterol (LDL-C); and HDL-cholesterol (HDL-C) and **E**, Triglycerides (TG) were determined in the plasma from fasted 5-months old rabbits of the indicated genotypes. For TC, HDL-C and TG: WT, n=24; *IDOL*^+/-^, n=22; *IDOL*^-/-^, n=25. For LDL-C: WT, n=21; *IDOL*^+/-^, n=18; *IDOL*^-/-^, n=19. *p<0.05 and **p<0.01. All data are mean±SEM. (A, B, C) Student’s t-test, (E, F) Kruskal-Wallis test and Dunn’s multiple comparison test for TC, LDL-C and TG. ANOVA and Tukey’s multiple comparison test for HDL-C. *p<0.05, **p<0.01.

### Rabbits as a model to study IDOL function

To further characterize the IDOL function in the rabbit liver, with the ultimate objective of providing a model feasible for preclinical studies involving IDOL inhibition, we generated an IDOL KO rabbit using CRIPR/Cas9 technology (Figure S1A-C) which introduced a 22-bp deletion resulting in a frameshift and loss of *IDOL* mRNA stability, likely from nonsense-mediated mRNA decay.^54^ Functionality of the deletion was determined using ear fibroblasts isolated humanely from *IDOL*^-/-^ and WT littermate controls on standard diet (Figure S1D). LDLR and ApoER2 were increased 4- and 2.5 fold, respectively. VLDLR was not detectable in the WT but the VLDLR Type I (full length) and Type II (lacking the O-linked sugar region) isoforms were readily detectable in the *IDOL*^-/-^. No appreciable changes were evident in LRP1 (LDL Receptor Related Protein 1), a protein involved in removal of atherogenic lipoproteins. Additionally, LDLR protein abundance was found to be 2.5-fold higher in the adrenal gland. These data are consistent with prior reports indicating increased LDLR protein in various tissues beyond liver in association with reduced *IDOL* expression.^28^ In the liver of *IDOL*^-/-^ rabbits on SD there was approximately a 2.2-fold increase in LDLR abundance relative to the WT rabbits (Figure S1E and F). SREBP-2 activates *LDLR* and *PCSK9* transcription in response to low intracellular cholesterol and the PCSK9 protein promotes LDLR degradation in lysosomes to ensure homeostasis.^55^ Conversely, *IDOL* and the *ABCG1* and *ABCA1* transporters are LXRα-targets activated by rising intracellular cholesterol.^38^ We determined that loss of IDOL resulted in increased LDLR protein without appreciable changes in PCSK9 abundance (Figure S1F). There were no changes in the relative mRNA levels of *ABCA1* and *ABCG1* in the liver of the same rabbits (Figure S1G). These data suggest that there is no appreciable activation of LXR or SREBP transcriptional pathways in the liver of *IDOL*^-/-^ rabbits on SD and reinforces that increased LDLR abundance is likely the result of loss of IDOL activity.^56,57^

### Decreased baseline cholesterolemia in IDOL KO rabbits on a standard-diet

We next determined whether loss of IDOL and ensuing increase in LDLR could alter plasma lipid profiles in male and female wildtype (WT), and littermate heterozygous (*IDOL*^+/-^), and homozygous (*IDOL*^-/-^) knockout rabbits on SD. The TC, LDL-C and HDL-C showed an overall significant decrease in *IDOL*^-/-^ rabbits, and a trend towards decrease in *IDOL*^+/-^ rabbits (Figure 1D). Compared to WT littermates, total cholesterol was 12% and 33% lower in *IDOL*^+/-^ and *IDOL*^-/-^ rabbits, respectively, with stronger effects in the females (Figure S2A). The decrease in TC was due to reduced LDL-C (38% and 56%, *IDOL*^+/-^ and *IDOL*^-/-^, respectively, Figure 1D, Figure S2B) and HDL-C (19% and 30% decrease, Figure 1D, Figure S2C). On SD, the triglycerides (TG), remained unchanged in the three genotypes, irrespective of sex (Figure 1E, Figure S2D). These data indicate that basal levels of plasma cholesterol are reduced in IDOL KO rabbits.

### Reduced diet-induced hyperlipidemia in IDOL KO rabbits

To study the role of IDOL during hyperlipidemia and atherosclerosis, the rabbits were fed a HCD for 16 weeks. While the average TC increased rapidly to over 1000 mg/dL after 8 weeks on HCD, the average of TC in *IDOL*^-/-^ rabbits remained below 250 mg/dL for the duration of the treatment (225 ± 28 mg/dL at 16 weeks) (Figure 2A). The total burden of plasma cholesterol in the 16 weeks of HCD, represented by the area under the curve (TC-AUC), was 82% lower in *IDOL*^-/-^ rabbits than in WT (Figure 2B), with both males and females showing highly significant reduction in the respective AUCs (Figure S3A-B). The *IDOL*^+/-^ rabbits showed a delay in development of hyperlipidemia resulting in a moderate (27% lower than WT) TC burden. Paralleling those data, while WT rabbits on HCD steadily increased their TG levels by 2.7-fold in average (31.2±1.6 mg/dL to 84.1±12.5 mg/dL), *IDOL*^-/-^ rabbits showed significantly lower TG for the 16 weeks of HCD-feeding (Figure 2C and D), *de facto* remaining within the range observed on SD for both genotypes (31.2 ±1.6 and 31.37±1.8, for WT and *IDOL*^-/-^, respectively). Meanwhile, the heterozygous showed delayed increase and a trend to intermediate AUC levels. These data indicate efficiently blocked HCD-induced hypertriglyceridemia by *IDOL*^-/-^, while simultaneously reducing the TC burden.

**Figure 2.**
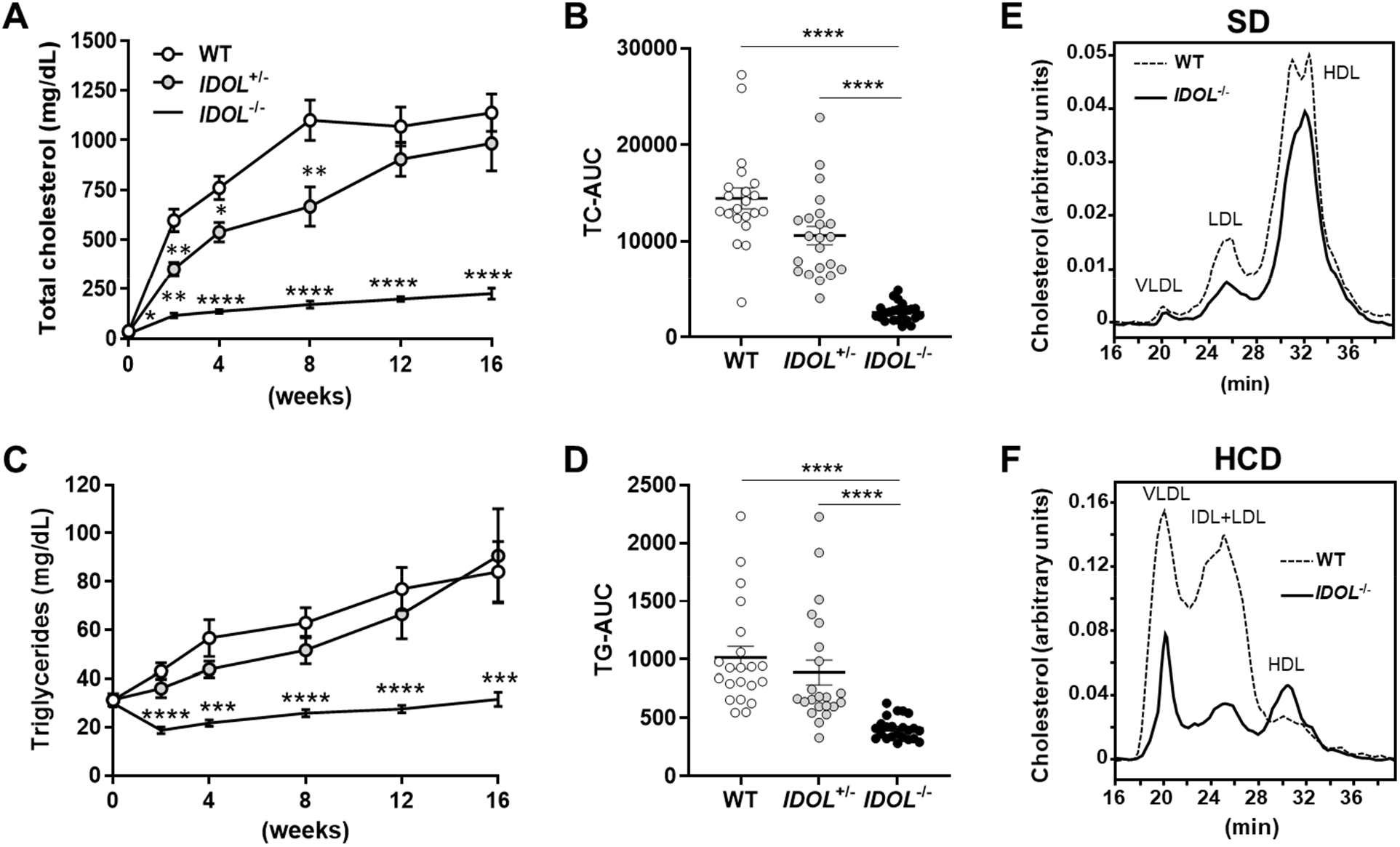
Reduced diet-induced hyperlipidemia in IDOL KO rabbits. Five months old rabbits, both sexes, wild type (WT) and *IDOL*^+/-^ and *IDOL*^-/-^ littermates on SD were switched to a HCD for 16 weeks. At the indicated time points, plasma was evaluated for **A**, Total cholesterol, with **B**, corresponding areas under the curves (TC-AUC). **C**, Triglycerides, with **D**, TG-AUC. WT, n=21; *IDOL*^+/-^, n=22; and *IDOL*^-/-^, n=23 (both sexes combined). **E** and **F**, Representative FPLC chromatogram of lipoprotein fractions from fasted male rabbits on SD (E) or after 16 weeks on HCD (F). All data are shown as mean± SEM. (A, B) RM two-way ANOVA with Geisser-Geenhouse correction and Dunnett’s multiple comparisons test. (C, D) Kruskal-Wallis test with Dunn’s multiple comparisons test. * p<0.05; ** p<0.01; *** p<0.001; ****p<0.0001.

Accordingly, the lipoproteins separated by FPLC from plasma of rabbits fed a SD showed differences in cholesterol contents only in the LDL and HDL particle traces (Figure 2E). Conversely, after 16 weeks on HCD, the FPLC analysis showed marked decreased cholesterol in all elution fractions (VLDL, ILD+LDL, Figure 2F). Further characterization by ultracentrifugation of plasma from WT and *IDOL*^-/-^ rabbits showed that cholesterol was reduced in all ApoB lipoproteins by 86%, 83%, 85%, and 79% in VLDL, IDL, LDL, and small LDL, respectively (Figure S3C). TG was reduced in the ApoB fractions by 69%, 68%, 67%, and 54% in VLDL, IDL, LDL, and small LDL, respectively in the *IDOL*^-/-^ (Figure S3D). When broken down by sex, both males and females had reduced cholesterol in the ApoB fractions, while females showed significant cholesterol increase in the ApoA fractions (Figure S3E and F). Collectively, these findings highlight a primary role for IDOL in TC and TG homeostasis in HCD.

### IDOL KO inhibits atherogenesis in the aorta and coronary artery

These findings indicate that interfering with IDOL activity could effectively reduce two main drivers of atherosclerosis, TC and TG, simultaneously when in an atherogenic diet and suggest a protective effect in atherosclerosis for IDOL loss-of-function. To assess this prediction, atherosclerosis was evaluated in the aorta and coronary arteries after 16 weeks on HCD in WT and *IDOL*^-/-^ rabbits. Atherosclerotic lesions, evidenced by *en face* staining with Sudan IV, were significantly reduced in the aorta of *IDOL*^-/-^ rabbits to less than 10% of the WT and were virtually confined to the aortic arch area (Figure 3A and B, Figure S4A). Histopathology of the aortic arch area showed thin macrophage-rich early lesions in *IDOL*^-/-^ rabbits compared to the thick fibroatheroma (with fibrous cap and necrotic core) present in the same area in WT rabbits (Figure S4B). In both sexes, *IDOL*^-/-^ showed significant reduction in atherosclerosis lesions throughout the whole aorta (Figure S4C and D). When compared to WT, *IDOL* ^+/-^ rabbits also showed a trend to decreased lesion areas in both sexes, albeit without achieving statistical significance overall, or in the combined data for both sexes (not shown). Analysis of the atherosclerotic lesions in the origin of the left coronary artery uncovered that the lesion area was 73% smaller in *IDOL* ^-/-^ than in WT rabbits (Figure 3C and D), with highly significant differences independent of sex (Figure S4E and F). Meanwhile, development of coronary atherosclerosis in *IDOL*^+/-^ rabbits from either sex (Figure S4E and F) or in the two sexes combined (not shown) was indistinguishable from the WT. Overall, these findings indicate that *IDOL* ^-/-^ rabbits are remarkably protected against atherosclerosis development in HCD-feeding, associated with reduced hyperlipidemia and independent of sex.

**Figure 3.**
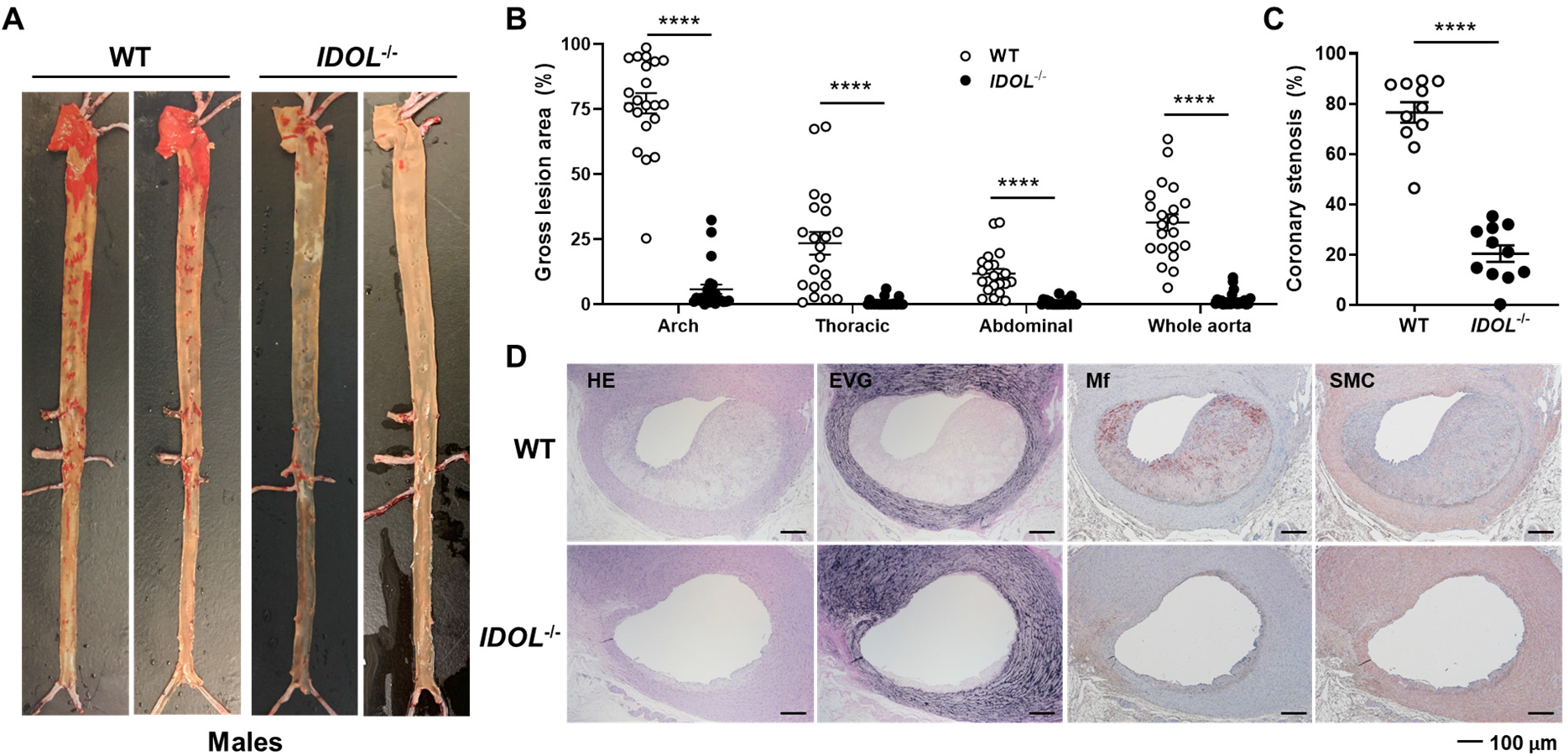
IDOL KO inhibits atherogenesis in the aorta and coronary artery. **A**. Representative pictures of aortic atherosclerosis in WT and *IDOL*^-/-^ male rabbits after 16 weeks on HCD. Lipids in the arterial wall were stained *en face* with Sudan IV to evidence the atherosclerotic lesions. **B**. Atherosclerotic lesions in 3 regions of the aorta (arch, thoracic and abdominal) and in the whole aorta were measured and expressed as the percentage (%) of red-stained area in the total area of each region. WT, n=21 (M/F=11/10); *IDOL*^-/-^, n=23 (M/F=9/14). **C**, Dot plot showing coronary atherosclerosis expressed as the percentage of coronary stenosis (%) calculated from the area of coronary atherosclerotic lesions relative to the area of the arterial lumen (n=11, each genotype, M/F= 6/5). Scale bar: 100 μm. **D**, Representative pictures of coronary atherosclerosis in WT and *IDOL*^-/-^ male rabbits. Samples were stained with hematoxylin and eosin (HE) and Elastica van Gieson (EVG). Macrophages (Mf) and smooth muscle cells (SMC) were immunostained with RAM11 and HHF35 antibodies, respectively. Scale bar, 100μm. All data are shown as mean±SEM. (B) Mann-Whitney U-test. (C) Unpaired t-test. **** p<0.0001.

### Steatosis was not exacerbated in the liver of IDOL KO rabbits on cholesterol-rich diet

Increase of IDOL targets in tissues in the *IDOL*^-/-^ rabbits can enhance uptake of circulating lipids and contribute to lower TC and TG levels in their plasma. In the liver, increased uptake of lipids creates potential for steatosis, from activation of hepatic LXR in the rabbit, unlike in mice.^58^ Therefore, we addressed the liver gross morphology and measured the cholesterol and triglycerides contents in the liver at endpoint of HCD. There were no differences in liver weight, or liver weight normalized by body weight (Figure 4A). The cholesterol or triglycerides per gram of liver tissue were indistinguishable between WT and *IDOL*^-/-^ rabbits (Figure 4B). The liver enzyme alanine transaminase (ALT) remained unchanged and aspartate transaminase (AST) was significantly reduced in *IDOL*^*-/-*^, indicating unlikely liver damage (Figure 4C). Histology indicated no obvious differences in the liver between WT and *IDOL*^-/-^ on HCD (Figure 4D). These data suggest that loss of IDOL can protect the rabbits against HCD-induced atherosclerosis without promoting liver steatosis.

**Figure 4.**
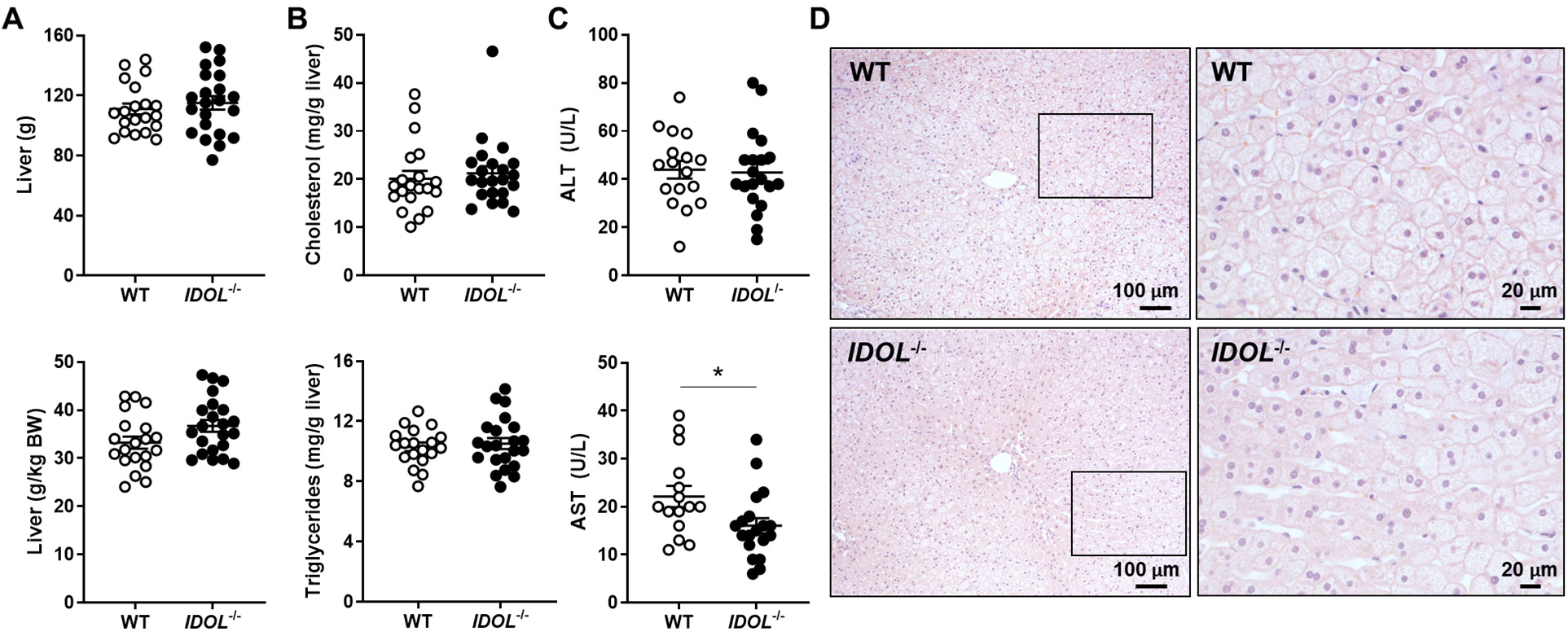
Steatosis was not exacerbated in the liver of IDOL KO rabbits. Livers were harvested from rabbits after 16 weeks on HCD. **A**, Liver weight (top) and liver weight normalized by total body weight (g/kg BW, bottom). **B**, Cholesterol (top) and triglycerides (bottom) contents in the liver (mg/g of liver). A and B, WT, n=20 (M/F=10/10); *IDOL*^-/-^, n=22 (M/F=8/14). **C**, Alanine aminotransferase, ALT, in WT (n=17; M/F=9/8) and *IDOL*^-/-^ (n=20; M/F=7/13) (top); and Aspartate aminotransferase, AST, in WT (n=15; M/F=7/8) and *IDOL*^-/-^ (n=19; M/F=6/13) (bottom), as determined in the serum of rabbits at 12 weeks on HCD. **D**, Representative images of liver histology samples stained by hematoxylin & eosin. Scale bar, 100μm. The areas indicated by the rectangles are shown at higher magnification in the pictures on the right. Scale bar, 20μm. All data are shown as mean±SEM. (A, Triglycerides in B) unpaired Student’s t-test. (Cholesterol in B) Mann-Whitney U-test. (C) Unpaired Student’s t-test, *p<0.05.

### Preserved TG clearance in IDOL KO rabbits on HCD

TC levels were only moderately increased while TG was not increased in average in plasma of *IDOL* ^-/-^ rabbits after 16 weeks on HCD when compared to SD (Figure 5A and B). Additionally, TG was reduced in the pro-atherogenic low density lipoprotein fractions (Figure S3). To investigate whether enhanced TG clearance in *IDOL* ^-/-^ rabbits could contribute to this phenotype, we performed intravenous fat tolerance test with Intralipid emulsion. We found that HFD significantly reduced the TG clearance in the WT rabbits in HCD while had no effect on the IDOL-/- rabbits, which preserved efficient TG clearance, comparable to those in SD (Figure 5C and 5D). Lipoprotein lipase (LPL) plays a crucial role in the plasma metabolism of VLDL and chylomicrons,^59^ the fractions most significantly reduced in TG contents in *IDOL* ^-/-^ rabbits (Figure S3D). Therefore, we assessed LPL activity as we previously described.^60^ While no differences were detected in SD, there was a significant increase in LPL activity in both genotypes in HCD with significantly higher levels in the *IDOL* ^-/-^ rabbits (75% in *IDOL* ^-/-^ vs 38% in WT). Collectively, these data indicate that, at least in part, IDOL KO averts the adverse effects of HCD through increased TG clearance and enhanced LPL activity.

**Figure 5.**
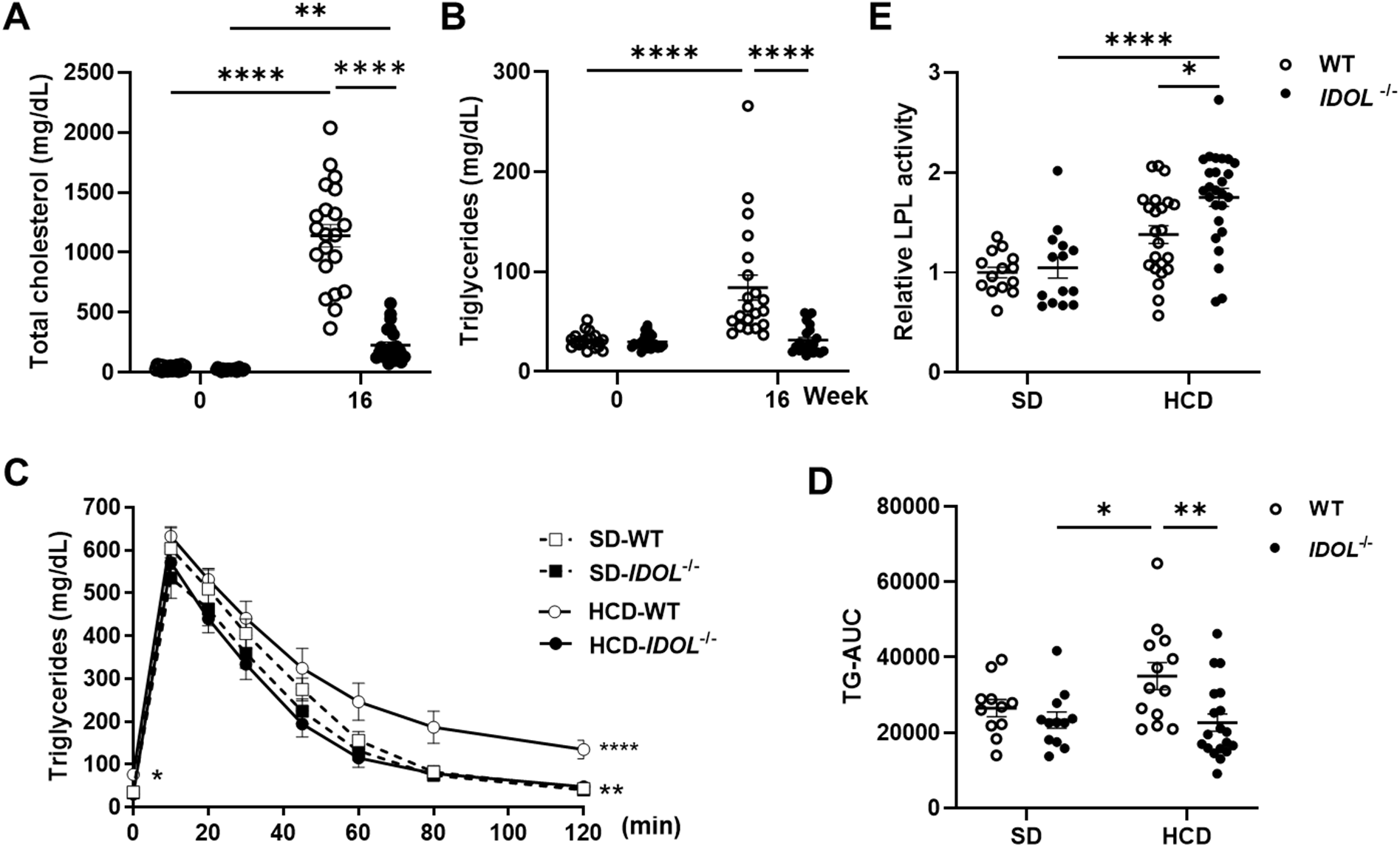
Preserved TG clearance in IDOL KO rabbits on HCD. **A**, Total Cholesterol and **B**, Total triglycerides in plasma at 16 weeks on HCD. WT, n=19; *IDOL*^-/-^, n =21. **C**, TG clearance using Intralipid emulsion test (2ml/kg of body weight) in rabbits on SD (dash lines) or on HCD (solid lines). WT, n=13 (M/F, 7/6); *IDOL*^-/-^, n=19 (M/F, 8/11). **D**, Area under the curves for the graph in C. **E**, Post-heparin (30U/kg of body weight) LPL activity assay. WT-SD, n=14; *IDOL*^-/-^-SD, n=14; WT-HCD, n=23; *IDOL*^-/-^-HCD, n=26. All data are shown as mean±SEM. (A, B) Mann-Whitney U-test. (C) Two-way ANOVA with Sidak’s multiple comparison test. *p<0.05 (WT vs KO in SD); **p<0.01 (WT vs KO in HCD); ****p<0.0001 (SD vs HCD for WT). (D) Two-way ANOVA with Tukey’s multiple comparison test. (E) Two-way ANOVA with Sidak’s multiple comparisons test. * p<0.05, **p<0.01, ****p<0.0001.

### Increased secondary bile acids in the feces from IDOL KO rabbits

IDOL KO affords protection against atherosclerosis without steatosis. To perform an initial characterization of the handling of excess cholesterol by the liver in the IDOL KO model, we focused on three main pathways: cholesterol efflux, cholesterol secretion into bile and secretion of bile acids. The ATP binding cassette subfamily A member 1 (ABCA1) and subfamily G member 1 (ABCG1) promote efflux of both neutral lipids and phospholipids from hepatocytes and macrophages.^57^ *ABCA1* and *ABCG1* are regulated by LXR and ERα.^61^ After 16 weeks of HCD, females presented significant upregulation of *ABCA1* and *ABCG1* expression by 1.9- and 2.0-fold, respectively in the liver of *IDOL*^-/-^ rabbits (Figure S5). Meanwhile, their relative expression in the adrenal gland, an organ with abundant LDLR, appeared to be independent of genotype. These data indicate that, in females, enhanced cholesterol efflux may contribute to preventing steatosis in the liver of the *IDOL*^-/-^ rabbits when on HCD.

Next, we addressed the presence of cholesterol and biliary acids in feces. HCD-feeding resulted in daily feces being decreased in both WT and *IDOL*^-/-^ rabbits by 25% and 28%, respectively, with no differences between the two genotypes within each diet (Figure 6A) or by sex (Figure S6A and B). Although daily cholesterol excretion was significantly increased on HCD, there were no differences across genotypes for each diet (Figure 6B). HCD feeding increased cholesterol excretion in feces per day in both WT and *IDOL*^-/-^ rabbits by 3.2- and 3.3-fold, respectively, with no differences by sex (Figure S6C and D). These data suggest that excretion of cholesterol in the bile may not contribute significantly to preserve lipid homeostasis in the IDOL KO livers.

**Figure 6.**
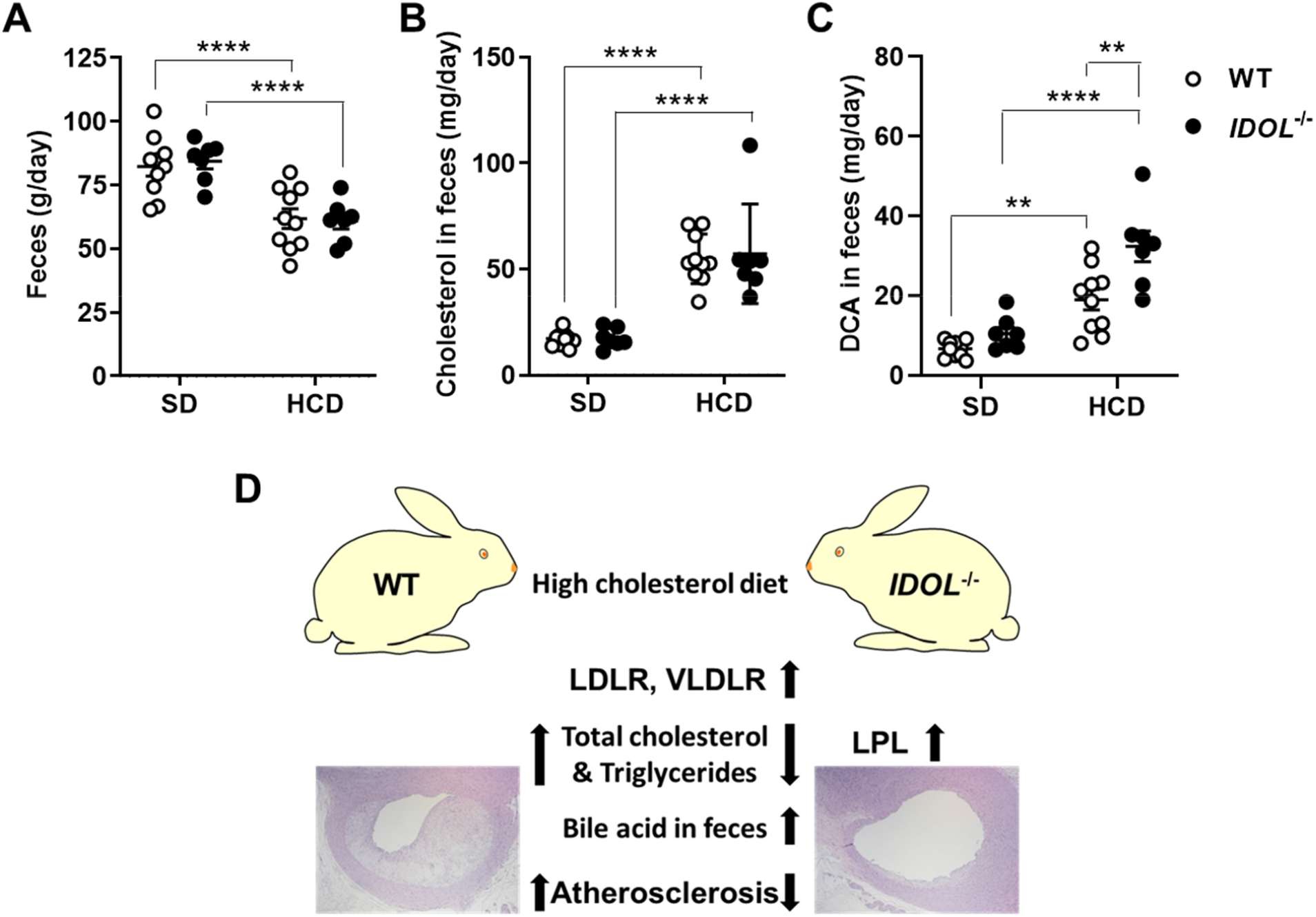
Increased bile acid excretion in the IDOL KO rabbits. **A**, Average daily weight of feces (WT, n=10; *IDOL*^-/-,^ n=7, sexes combined). **B**, Average daily cholesterol in feces. **C**, Deoxycholic acid (DCA) in feces. **D. Graphical summary of the findings**. All data are shown as mean±SEM. (A, B) Two-way ANOVA with Sidak’s multiple comparisons test. (C) Two-way ANOVA with Tukey’s multiple comparisons test. **p<0.01, ****p<0.0001

Finally, we analyzed the metabolites of cholesterol as bile acids by mass spectrometry. Of 18 bile acids analyzed (Supplemental Table 1), we were able to quantify 15 of them (Figure S6E and F). Deoxycholic acid (DCA) and lithocholic acid (LCA), were the two major bile acids in feces, as previously reported.^62,63^ Specifically, when on HCD, DCA was increased in both genotypes with a 1.7-fold further increase in the *IDOL*^*-/-*^ (Figure 6C). In fact, although it did not achieve statistical significance in the grouped analysis, *IDOL*^-/-^ showed a 1.5-fold increase of DCA in feces on the SD as well. We found no significant differences in LCA (Figure S6E and F). Collectively, these data indicate that the bile acid pathway is enhanced on HCD and that this effect is more pronounced in the absence of *IDOL*, likely contributing to preventing excess hepatic lipid accumulation in the *IDOL*^-/-^ rabbits.

## DISCUSSION

To help accelerate progress towards potential IDOL-based therapies, we introduce here a rabbit IDOL KO model created using CRISPR/Cas9 and provide an initial characterization in the context of atherosclerosis. We offer proof-of-concept of preservation of the LXR/IDOL/LDLR axis in the liver of rabbits. We found that *IDOL* expression in the liver was under LXR-regulation and is increased in response to HCD. On SD, loss of IDOL increased LDLR, ApoER2 and VLDLR stability. *IDOL*^*-/-*^ rabbits displayed reduced TC, LDL-C, HDL-C. On HCD, IDOL KO resulted in a simultaneous reduction of TC and TG in plasma, associated with enhanced TG clearance and LPL activity in plasma and increased bile acid excretion in feces. Consequently, IDOL KO protected rabbits against atherosclerosis on HCD, remarkably without apparent effects on liver steatosis.

Dyslipidemia and sustained high lipid burden are the main drivers of atherosclerosis. Combination therapies, albeit effective in reducing total cholesterol, are restricted in their efficacy due to physiological feedback loops and the lingering levels of triglycerides. Clinical trials of LXR agonists in humans were stopped because of increased hepatic steatosis and dyslipidemia.^25, 26^ Over the last 15 years, human genetic studies^41,42,64-66^ and mechanistic insights^35-37^ posit IDOL inhibition as an attractive approach for the treatment of hyperlipidemia, although differential LXR-dependent regulation of hepatic *IDOL* in mice and primates has slowed progress towards that goal.^29^ In mice, *Idol* has low impact on plasma LDL, likely due to limited sterol-dependent regulation of *Idol* expression in their liver and there is no appreciable upregulation of LDLR in the liver of *Idol*^*-/-*^ mice. Additionally, mice do not develop atherosclerosis in the absence of genetic manipulation. Recently, humanized liver models showed LXR-dependent *IDOL* activation, albeit without evidence of atherosclerosis development either.^67,68^ In contrast, in non-human primates, LXR activation induces hepatic IDOL resulting in reduced LDLR protein and increased plasma LDL-C. Conversely, IDOL knockdown by antisense oligonucleotides reduces the effects of the LXR agonist on LDL-C levels. This not only uncovered species-specific differences but also underscores the contribution of liver LDLR to plasma lipid clearance.^69^ Compared with primate and pig models, more expensive and less amenable to genetic manipulation, rabbit is an established model to study diet-induced atherosclerosis and associated lipoprotein metabolism, with features relevant to human disease.^47,70^

One notable finding is that, on HCD, IDOL KO can prevent increases in plasma TC and TG on the atherogenic diet. Mendelian randomization analyses showed that LPL variants associated with lower TG reduced the risk of coronary heart disease, to a level comparable to LDLR variants lowering LDL-C, indicating that concurrent reductions in LDL-C and TG could have enhanced benefits for CVD.^71^ We found that IDOL KO resulted in a very significant reduction of TC and TG in the ApoB/VLDL fractions, the predominant atherogenic agents. TG-rich lipoproteins are an atherogenic risk factor.^11,12^ Reduced plasma TG in IDOL KO may be driven by increased VLDLR-dependent uptake or reduced lipogenesis. Additionally, LPL contributes to normal lipoprotein metabolism and energy balance,^72^ and is the rate-limiting enzyme for TG hydrolysis from chylomicrons and VLDL.^59^ On SD, we found no differences in TG levels or LPL activity between WT and IDOL KO. On HCD, *IDOL*^-/-^ abrogated TG increase in plasma and preserved high TG clearance, comparable to that on SD. This was associated with increased LPL activity beyond that observed for the WT. The improved phenotype in the IDOL KO model may be reflecting altered interactions of LPL with its regulators, including ApoCIII or ANGPTL3 (inhibitors) or ApoCI (activator) produced in the liver. We recently reported that rabbit ApoCIII KO increased LPL activity with a remarkable reduction in atherosclerosis concomitant to reduced β-VLDL, and blunting of TC and TG increases.^60^ The IDOL KO rabbit model will provide a unique resource to address whether IDOL KO affects the production or secretion of LPL or of its regulators or whether it alters the composition its lipoprotein substrates. Of potential clinical significance, IDOL KO did not increase steatosis in HCD, suggesting improved handling of excess lipids in the liver. Three main pathways contribute to eliminate excess cholesterol from the liver: increased secretion of cholesterol, bile acid secretion in feces or efflux mediated by ABCA1 and ABCG1.

Cholesterol contents in the feces did not differ between IDOL KO and WT rabbits on either diet. On the other hand, the increase of DCA in feces in HCD-feeding was significantly higher in the IDOL KO rabbits, indicating that altered bile acid metabolism may be a main factor in counteracting the increased influx of cholesterol in the lDOL KO liver. Whether the increased DCA in feces, a secondary biliary acid produced by the gut microbiome from DCDA, is the result of increased output from the liver of the precursor, reduced DCDA uptake to the enterohepatic circulation by the intestine or microbiome remodeling,^73^ will require extensive research in this model beyond the scope of this initial characterization.

Although both sexes showed positive and equivalent outcomes from IDOL KO in dyslipidemia and atherosclerosis, we uncovered sex-differences in our data reminiscent of observations in humans.^74,75^ *ABCA1* and *ABCG1*, encode two key enzymes in reverse cholesterol transport and are known targets of both LXR and ERα.^61^ ABCG1 mediates efflux to HDL, whereas ABCA1 is required for efflux to ApoA1, in addition to being one rate-limiting step in the formation of HDL. *ABCA1* and *ABCG1* were increased in the liver in female *IDOL*^-/-^ rabbits concurrent with significantly increased cholesterol in the HDL/ApoA1 lipoprotein fractions in plasma. It is possible that both cholesterol efflux and biliary acid secretion contribute to prevent liver steatosis in the IDOL KO rabbits, although their relative contribution and regulation remain to be determined. Studies addressing such sex differences may have profound relevance to development of IDOL-based therapies and may be facilitated by this rabbit model. Additionally, future studies should address the fundamental question on whether and how IDOL KO might uncouple some of the delicately regulated interactions involving LXR, SREBP, PPARα (amongst other factors) and their coordinated functions to ensure lipid homeostasis in the liver in the proatherogenic diet.

In closing, rabbit models have helped make significant contributions of clinical relevance to CVD and beyond. It is our hope that this initial characterization of the IDOL KO rabbits will provide the scientific community with an adequate framework and a valuable resource to continue deepening our understanding of the LXR/IDOL dynamics in the liver and the physiologic consequences of targeting IDOL to reduce dyslipidemia and atherosclerosis.

## Nonstandard Abbreviations and Acronyms

ABCA1: ATP binding cassette subfamily A member 1
ABCG1: ATP binding cassette subfamily G member 1
ALT: alanine transaminase
ApoER2: apolipoprotein E receptor 2
AST: aspartate transaminase
BW: body weight
CETP: cholesteryl ester transfer protein
CRISPR: clustered regularly interspaced short palindromic repeats
CRP: C-reactive protein
CVD: cardiovascular disease
DCA: deoxycholic acid
FPLC: fast-performance liquid chromatography
HCD: high-cholesterol diet
HDL: high-density lipoprotein
IDL: intermediate-density lipoprotein
IDOL: inducible degrader of theLDLR
KO: knockout
LCA: lithocholic acid
LDL: low-density lipoprotein
LDLR: LDL receptor
LPL: lipoprotein lipase
LXR: liver X receptor
oxLDL: oxidized low-density lipoprotein
qPCR: quantitative real time polymerase chain reaction
PCSK9: proprotein convertase subtilisin/kexin type 9
RXR: retinoid X receptor
SD: standard diet
SMC: smooth muscle cell
SREBP: sterol regulatory element binding transcription factor 2
TC: total cholesterol
TG: triglycerides
TRL: triglyceride-rich lipoprotein
VLDLR: very-low-density lipoprotein receptor
WT: wild type

## Sources of Funding

This work was supported by the National Institutes of Health grants HL147527, HL159900, HL109946 (Y.E. Chen), HL138139 and HL153710 (J. Zhang), and HL150233 (O. Rom). We utilized the services from the University of Michigan In-Vivo Animal Core supported by grant U2CDK110768).

## Disclosure

None

## HIGHLIGHTS

- The LXR/IDOL/LDLR axis is conserved in the rabbit liver and, unlike in mice, rabbit hepatic IDOL is upregulated on high cholesterol diet.
- IDOL deficiency in rabbits increased LDLR in the liver in association with reduced plasma total cholesterol on standard diet.
- In rabbits on high cholesterol diet, IDOL KO concurrently inhibited cholesterol increases and abrogated hypertriglyceridemia through enhanced lipoprotein metabolism and heightened bile acid excretion in feces.
- Consequently, IDOL KO protected rabbits against diet-induced atherosclerosis independent of sex and without aggravation of liver steatosis.
- Availability and further characterization of this animal model will accelerate clinical translation of IDOL inhibition as a potential strategy to further reduce the burden of hyperlipidemia in cardiovascular diseases.

## Notes

### Competing Interest Statement

The authors have declared no competing interest.

