## supplemental for "IDOL deficiency inhibits cholesterol-rich diet-induced atherosclerosis in rabbits"

1 Department of Internal Medicine, Frankel Cardiovascular Center; Center for Advanced Models for Translational Sciences and Therapeutics. University of Michigan Medical Center, Ann Arbor, MI 48109

2 Department of Pharmaceutical Sciences, College of Pharmacy, University of Michigan, Ann Arbor, MI 48109

3 Department of Pathology and Translational Pathobiology; Center for Cardiovascular Diseases and Sciences, Louisiana State University Health Sciences Center-Shreveport, Shreveport, LA, 71103

4 Division of Cardiac Surgery, Department of Surgery, Davis Heart and Lung Research Institute, The Ohio State University Wexner Medical Center, Columbus, OH, 43210

5 Department of Molecular Pathology, Faculty of Medicine, Graduate School of Interdisciplinary Research, University of Yamanashi, Yamanashi, Japan.

**Short title:** IDOL knockout inhibits atherosclerosis in rabbits

**Corresponding Authors**:

Y. Eugene Chen, MD, PhD, Department of Internal Medicine, Frankel Cardiovascular Center, University of Michigan Medical Center. 2800 Plymouth Rd, Ann Arbor, MI 48109,

Jifeng Zhang, PhD, Department of Internal Medicine, Frankel Cardiovascular Center, University of Michigan Medical Center. 2800 Plymouth Rd, Ann Arbor, MI 48109,

Minerva T. Garcia-Barrio, PhD, Department of Internal Medicine, Frankel Cardiovascular Center, University of Michigan Medical Center. 2800 Plymouth Rd, Ann Arbor, MI 48109,

**Generation of IDOL KO rabbits and husbandry**

New Zealand White rabbits (Covance Inc. Princeton, NJ) were used for this study. The IDOL knockout (IDOL KO, for the model; IDOL$^{-/-}$, for the genotype) rabbits were generated by CRISPR/Cas9 endonucleases as described previously.[1] In brief, based on the rabbit *IDOL* genomic sequence (Ensembl: ENSOCUG00000008005), we designed the guide RNA (gRNA, GGATCTCCCAGCAGATGGACggg) using the online software (http://crispor.tefor.net/crispor.py), which targets exon 2 (Supplemental Fig. 1A). We injected 53 embryos with Cas9/gRNA mix and transferred them to foster mothers. Five kits were born, all positive for IDOL gene targeting. The founders were bred with wild type rabbits to confirm germline transmission. From the F1 generation rabbits, we choose one with a 22bp deletion (AGCAGATGGACGGGCTGGCCCC) in exon 2 to establish the IDOL KO rabbit line used for the current study. The 22bp deletion after codon 63 introduced a frameshift, as determined by Sanger sequencing (Supplemental Fig. 1B), with 82 unrelated codons out of frame before the premature stop, and the consequent loss of the *IDOL* mRNA across all tissues tested (Supplemental Fig. 1C), likely due to early translation termination causing nonsense-mediated mRNA decay.[2]

All rabbits were housed individually in cages under constant 20°C temperature and 12-hour light/dark cycles. Rabbits were fed 120g/day of a standard diet (SD, Teklad global rabbit diet 2030, Envigo, Indianapolis, IN) or a high cholesterol diet (HCD, custom diet containing 0.3% cholesterol and 3% soybean oil added to the SD, Envigo) for 16 weeks (endpoint), which induces severe hypercholesterolemia (>500 mg/dL within a month) and atherosclerosis in the aorta of otherwise normal rabbits.[3] For acute lipid-loading experiments spanning 7 days and 6 weeks, the custom diet contained 1% or 0.6% cholesterol, respectively.[4,5] At endpoint, the rabbits were anesthetized with 4%–5% isoflurane inhalation and euthanized by exsanguination and thoracotomy.[6] All animal studies were performed in accordance with the animal protocols approved by the Institutional Animal Care and Use Committee (IACUC) at the University of Michigan. For initial characterization of *IDOL*$^{-/-}$ rabbits (Supplemental Figure 1, C-G)

**Induction of hyperlipidemia and atherosclerosis**

To induce hyperlipidemia and atherosclerosis, rabbits were fed 120g/day of the HCD described above for 16 weeks. The food intake was monitored daily. The body weights were recorded, and blood samples (2ml) were collected from the central auricular artery at 0, 2, 4, 6, 8, 12, and 16 weeks to determine total cholesterol (TC) and triglycerides (TG) as described below. Lipoprotein fractions were analyzed after 10 weeks on HCD as described below. At the endpoint, rabbits were fasted overnight and euthanized as described above.[6] Blood was washed out by saline perfusion through the left ventricle. Organs were dissected, weighed and samples were collected, processed and preserved in formalin or frozen in liquid nitrogen for further analyses.

**Analysis of plasma lipids**

TC, TG, and high density-cholesterol (HDL-C) in plasma were measured using commercially available assay kits (TC, cat# 990-02601, TG, cat# 290-63701, HDL-C, cat# 997-01301, Wako Diagnostics, Mountain View, CA) according to the manufacturer’s instructions. The low density-cholesterol (LDL-C) was analyzed by the Chemistry Laboratory of the Michigan Diabetes Research and Training Center (University of Michigan).

**Analysis of plasma lipoproteins**

Plasma lipoprotein fractions were isolated by sequential density ultracentrifugation of plasma from rabbits fed normal chow diet or at 10 weeks on HCD, as we described previously.[3] Seven main different density lipoproteins were isolated from each sample as follows: Density (g/mL) < 1.006, VLDL and chylomicron remnant; 1.006-1.02 (-1.02), IDL; 1.02-1.04 (-1.04), LDL; 1.04-1.06 (-1.06), small LDL and HDL1; 1.06-1.08 (-1.08) and 1.08-1.10 (-1.1), HDL2; and 1.10-1.21 (-1.21), HDL3. All fractions were dialyzed against phosphate-buffered saline (PBS) overnight. TC and TG levels in each fraction were measured as described above. Plasma lipoprotein profiles were also analyzed by fast-performance liquid chromatography (FPLC) as described previously.[7] The cholesterol in each of the 40 fractions collected was determined using the Cholesterol Fluorometric Assay Kit (Cayman Chemical, Ann Arbor, MI).

**Atherosclerotic lesion analysis in the aorta and left coronary artery**

Atherosclerotic lesions were quantified using the method we described previously.[3,8] The rabbit aortas were stained with Sudan IV for evaluation of gross atherosclerotic lesions and the measurements were done in a double-blinded manner. The lesion size was expressed as % of the lesion area relative to the total area of that region in each indicated section of the aorta, and the % area of all lesions relative to the total aorta. For histology analysis of the lesions, paraffin-embedded serial sections (5 μm) of the aortic arch area were stained by H&E and Elastica van Gieson (EVG). For immunohistochemistry, serial sections were stained with antibodies against macrophages (RAM11, Agilent Technologies, Inc., Santa Clara, CA) and smooth muscle actin (HHF35, Agilent Technologies, Inc., Santa Clara, CA).

Analysis of coronary lesions was performed as we recently reported.[9] In brief, the rabbit heart was sectioned into 5 blocks. Block 1 contains the main trunk of the left coronary artery which was sectioned and stained with H&E and immunostaining as above. The coronary stenosis was measured using ImageJ (3 sections/rabbit). Coronary atherosclerosis was expressed as the percentage of coronary stenosis (%) calculated from the area of coronary arterial atherosclerotic lesions divided by the area of the arterial lumen as previously reported.[3]

**Analysis of TG clearance**

The TG clearance test was performed as reported previously.[10] Briefly, rabbits on SD or after 14 weeks on HCD were fasted overnight and administered intravenously 2 ml/kg of body weight of Intralipid emulsion (Sigma-Aldrich, St. Louis, Mo). Blood samples were withdrawn at 0, 10, 20, 30, 45, 60, 80, 120 min after the emulsion was injected. TG levels in each blood sample were measured as described above.

**Plasma lipoprotein lipase activity**

Plasma lipoprotein lipase (LPL) activity was determined as we previously reported.[11] Briefly, rabbits (both males and females, on the indicated diet) were fasted for 16 hours, secured in a restrainer without anesthesia, and injected intravenously with heparin (30 U/kg), followed by blood collection after 10 min. LPL activity in post-heparin plasma was measured by an LPL activity assay kit (Cell Biolabs, Inc. San Diego, CA) according to the manufacturer’s instructions.

**Lipids in liver and hepatic pathology**

Lipids in tissue samples from the liver and adrenal were extracted with a chloroform free kit (Lipid extraction kit, # K216, BioVision, Milpitas, CA) according to the manufacturer's instructions. Cholesterol and triglyceride concentrations in the extracted lipids from these organs were measured as described above and expressed as mg/g of tissue. The paraffin-embedded liver samples were sliced (4 µm thick) and stained by hematoxylin and eosin (H&E) for the pathology evaluation. Liver enzymes in serum (ALT and AST) were determined by the In-Vivo Animal Core (IVAC) at the University of Michigan.

**Cholesterol and bile acids in feces**

The cholesterol excretion to the feces was addressed in an acute cholesterol loading experiment by feeding a modified HCD (HCD containing 1.0% cholesterol) for 1 week.[4] Three days before starting the HCD (baseline, rabbits on SD), and every day throughout the 7 days on HCD, feces were collected, and their weight recorded daily. To minimize daily variations, for each rabbit, feces from three days on SD and from days 5 to 7 on HCD were pooled by diet independently and stored at -80ºC for further analysis. The pooled feces were minced and mixed thoroughly before lipid and cholesterol extraction using the Folch method[12]. Cholesterol levels in the extracted lipids were measured by the Cholesterol Fluorometric Assay Kit (#10007640, Cayman Chemical, Ann Arbor, MI). Bile acids in the feces were extracted and analyzed by LC-MS/MS.[13] Briefly, 400 µl of 80% methanol/water solution were added per 100 µg of feces sample. The mixed solution was homogenized and centrifuged to collect the supernatant. Analysis was performed on 2 µl of each supernatant by LC-MS/MS at the Pharamacokinetic and Mass Spectometry Core at the University of Michigan core. Of 18 bile acids (Online Table 1), we were able to detect 15, with the secondary bile acids DCA and LCA being the most abundant, as previously described.[14]

**RNA isolation and quantitative real-time PCR (qPCR) analysis**

Total RNA from rabbit tissues were isolated with RNA Tissue Mini Kit (QIAGEN, 959034, Hilden, Germany) according to the manufacturer’s instructions. SuperScript™ III First-Strand Synthesis

System (Thermo Fisher Scientific, 18080051) and random primers were used to reverse transcribe RNA into cDNA. Gene expression was quantified by qPCR using iQ SYBR Green Supermix (1708882, Bio-Rad, Hercules, CA). The gene expression was normalized to the internal control β-actin or GAPDH. The primer sequences used are listed in Online Table 2.

**Protein extraction and Western blot**

Rabbit tissues (~100mg) were homogenized in T-PER™ tissue protein extraction reagent (ThermoFisher Scientific, 78510) supplemented with the cOmplete™ EDTA-free protease inhibitor cocktail (Roche, 11873580001, Penzberg, Germany) and PhosSTOP™ phosphatase inhibitor (Roche, 4906845001, Penzberg, Germany). Tissues were lysed at 4°C for 30min and centrifuged at 13,000 rpm for 15min to remove insoluble debris. Protein extracts were resolved in 10% SDS-PAGE gels and transferred to nitrocellulose membranes (BioRad, 1620115, Hercules, CA). The indicated primary antibodies were incubated at 4°C overnight and subsequently incubated with secondary antibody (1:10,000 dilution, Li-Cor Bioscience, Lincoln, NE) for 1 hour at room temperature. Bands were scanned using Odyssey Imaging System (Li-Cor Bioscience, Lincoln, NE) and quantified with the LI-COR Image Studio Software. For primary antibodies, please see Major Resources Table.

**Statistical analysis**

All data are expressed as mean±SEM. Statistical analyses were performed using GraphPad Prism 9, GraphPad Software, San Diego, CA. Statistical analyses for comparison between 2 groups or among 3 groups were selected based on the results of the Shapiro-Wilk test normality test for the distribution of each data set. For 2-group comparisons, two-tailed, unpaired Student's t-test (parametric) or Mann-Whitney U-test (nonparametric) were used. For ≥3-group comparisons, we performed ordinary one-way ANOVA and Tukey's multiple comparisons test (parametric) or Kruskal-Wallis and Dunn's multiple comparison test (nonparametric). Two-way ANOVA and Sidak's multiple comparison test were performed for the data obtained from multiple experimental time points or conditions. Repeated measures (RM) Two-way ANOVA was used for analysis of repeated by-weekly measures of lipid changes in plasma. Statistical significance was set at $P < 0.05$.

**Supplemental figures**

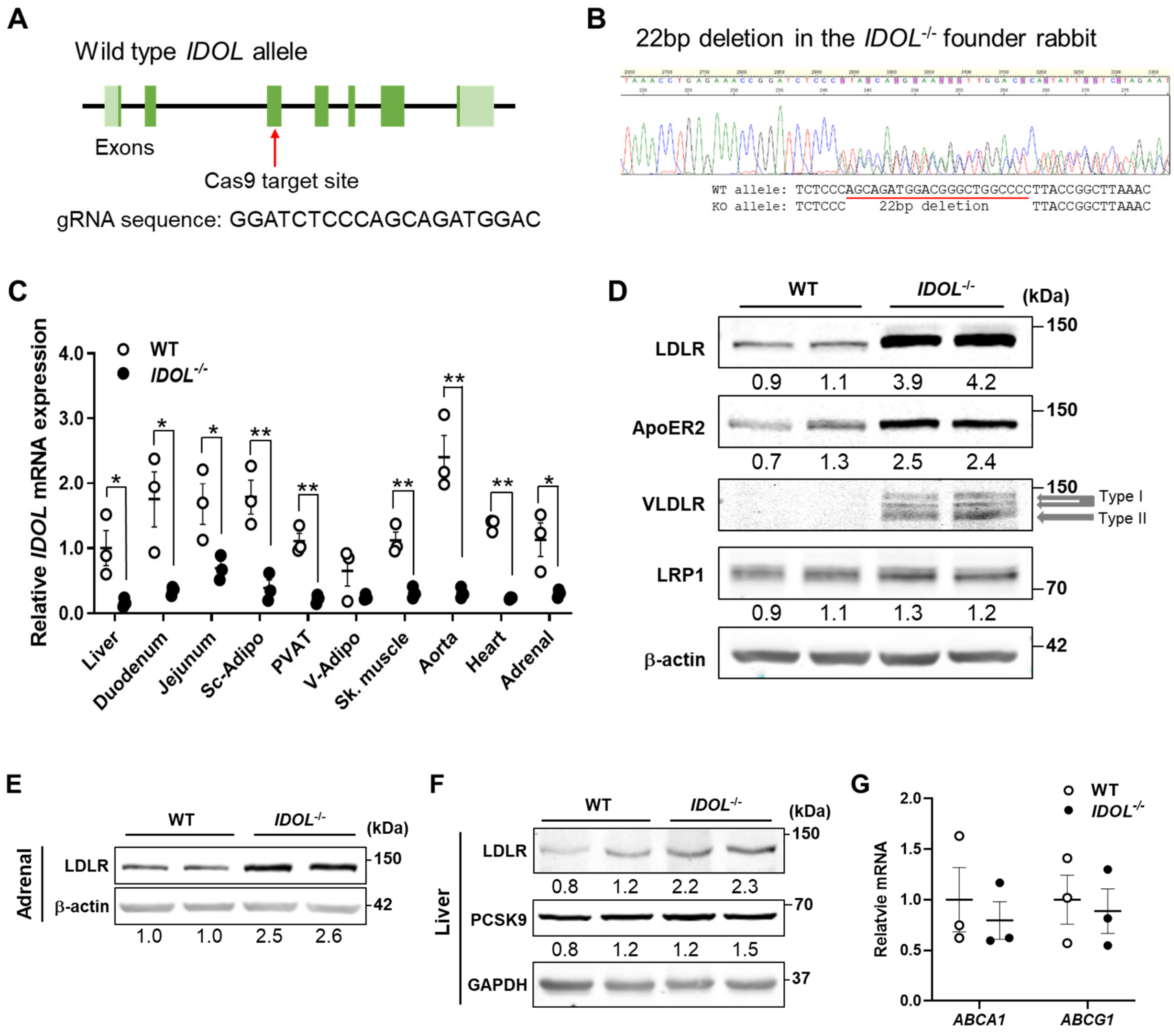


**Figure S1. Generation of IDOL KO rabbits and initial characterization**. **A**, Schematic representation of the strategy to generate *IDOL*$^{-/-}$ rabbits using CRISPR/Cas9, including the guide RNA sequence (gRNA). **B**, Verification of the 22bp deletion by Sanger sequencing. **C**, *IDOL* mRNA abundance, normalized by 18S, in the indicated tissues from wild type (WT) and *IDOL*$^{-/-}$ rabbits on standard diet (SD, 5-months old males, n=3, WT liver set to 1). Sc-Adipo, V-Adipo and PVAT, subcutaneous visceral and perivascular adipose tissue, respectively; Sk. muscle, skeletal muscle. **D**, Functional verification of *IDOL*$^{-/-}$ using ear fibroblasts humanely isolated from living WT and *IDOL*$^{-/-}$ rabbits and Western blot for the known IDOL targets (LDLR and ApoER2 and VLDLR) and the non-target LRP1 (LDL Receptor Related Protein 1). Relative abundance of each protein was determined by densitometry, normalized by β-actin, and is indicated under each lane. The WT average was set to 1. **E**, Relative abundance of LDLR protein in the adrenal gland of WT and *IDOL*$^{-/-}$ rabbits determined as in C. **F**, Relative abundance of PCSK9 and LDLR proteins in the liver of WT and *IDOL*$^{-/-}$ rabbits determined as in C, normalized by GAPDH . **G**, Relative mRNA expression of *ABCA1* and *ABCG1* in liver of the WT and *IDOL*$^{-/-}$ littermate rabbits on SD, with the WT set as 1, n=3. All data are shown as mean±SEM. (B) Student's t-test, independently comparing the two genotypes for each individual tissue. (G) Mann-Whitney test indicates p=0.7 (no significant differences). * $p<0.05$; ** $p<0.01$.

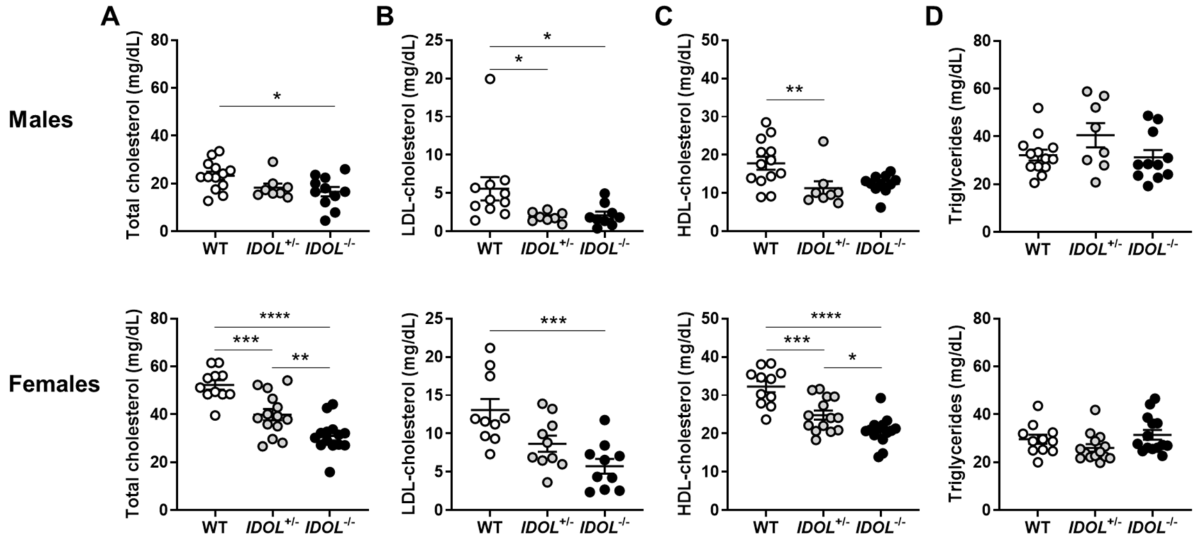


**Figure S2. Cholesterol and triglycerides by sex in IDOL KO rabbits on standard diet**. **A**, total cholesterol; **B**, LDL-cholesterol; **C**, HDL-cholesterol; **D**, Triglycerides in males (top panels) and females (bottom panels) was determined in plasma from 5-months old male and female rabbits of the indicated genotypes on a SD. A, C and D: WT, n=24 (male, 13; female, 11); *IDOL*$^{+/-}$, n=22 (male, 8; female, 14); *IDOL*$^{-/-}$, n=25 (male, 11, female, 14). B: WT, n=21 (male, 11; female, 10); *IDOL*$^{+/-}$, n=18 (male, 8; female, 10); *IDOL*$^{-/-}$, n=19 (male, 9, female, 10). Data are mean ± SEM. A, B and C males, D females: Kruskal-Wallis and Dunn's multiple comparisons test. A, B and C females, D males: ANOVA and Tukey's multiple comparison test. *$p<0.05$, **$p<0.01$, ***$p<0.001$ and ****$p<0.0001$.

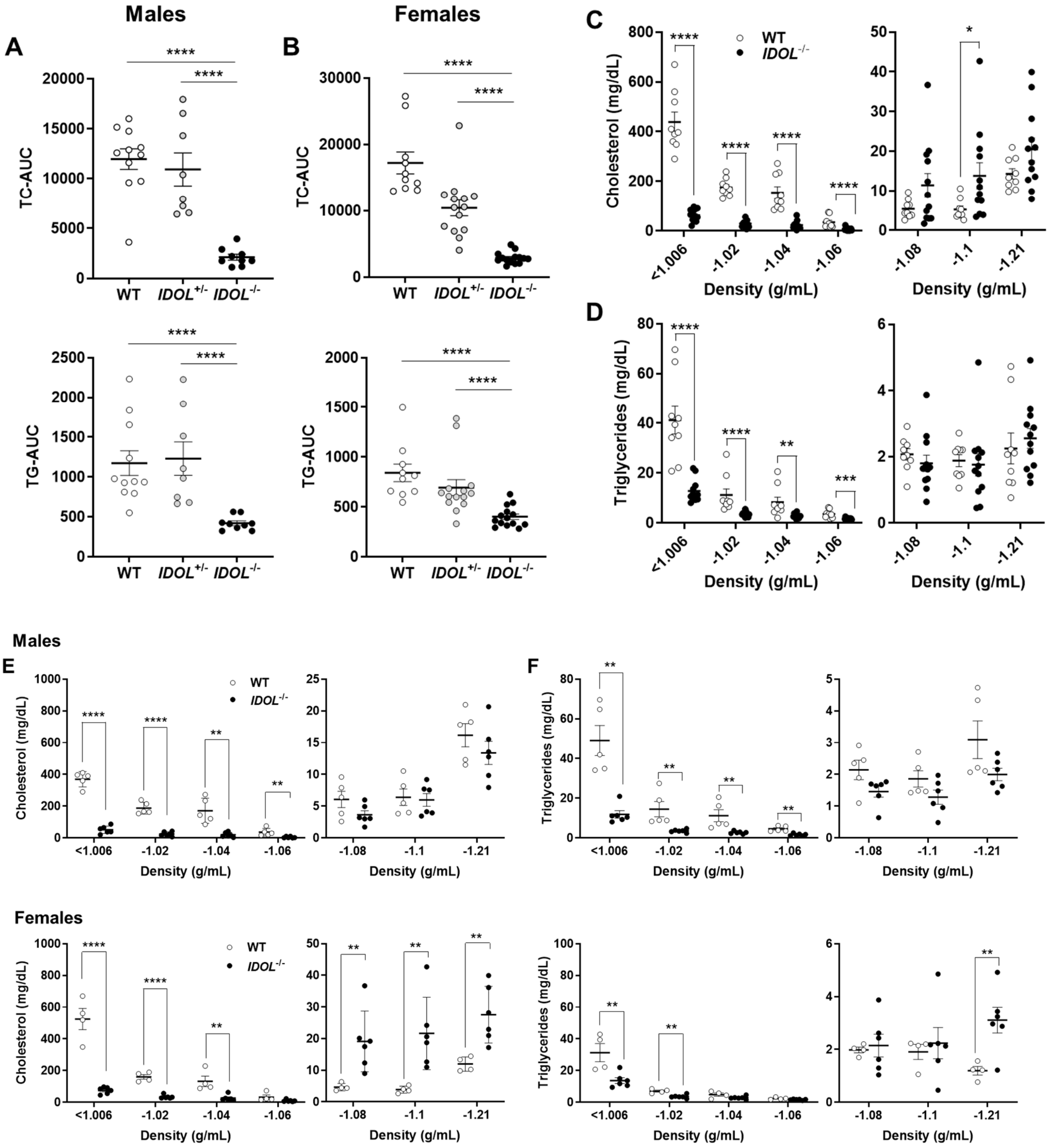


**Figure S3. Reduced diet-induced hyperlipidemia in IDOL KO rabbits is associated with changes in lipoprotein profiles.** Five months old rabbits, *IDOL*$^{+/-}$ and *IDOL*$^{-/-}$ and wild type (WT) and littermates, were fed a high cholesterol diet (HCD, 0.3% cholesterol and 3% soybean oil added to the 2030 Teklad global rabbit diet, Envigo) for 16 weeks. Data are areas under the curves (not shown) of total cholesterol (TC-AUC) or triglycerides (TG-AUC) from **A**, males and

**B**, females. WT, n=21 (male, 11; female, 10); *IDOL*$^{+/-}$, n=22 (male, 8; female, 14); and *IDOL*$^{-/-}$, n=23 (male, 9; female, 14). **C**, cholesterol, and **D**, triglycerides in lipoprotein fractions from sequential ultracentrifugation of plasma from fasted rabbits at 10 weeks on HCD. WT, n=9 (male, 5; female, 4); and *IDOL*$^{-/-}$, n=12 (male, 6; female, 6). **E** and **F**, data by sex from C and D above. Density (g/mL) < 1.006, VLDL and chylomicron remnant; 1.02 , IDL; 1.04, LDL; 1.06, small LDL and HDL1; 1.08 and 1.10, HDL2; and 1.21, HDL3. All data are mean±SEM. (A, B) Kruskal-Wallis test with Dunn's multiple comparisons test. (C) Student's t-test (for fractions 1.006, 1.02, 1.04, and 1.21); Mann-Whitney U-test (for fractions 1.06, 1.08, 1.10). (D) Student's t-test (for 1.06); and Mann-Whitney U-test for the rest of the fractions. Data are mean±SEM. *p=0.05, **p<0.01, ****p<0.0001.

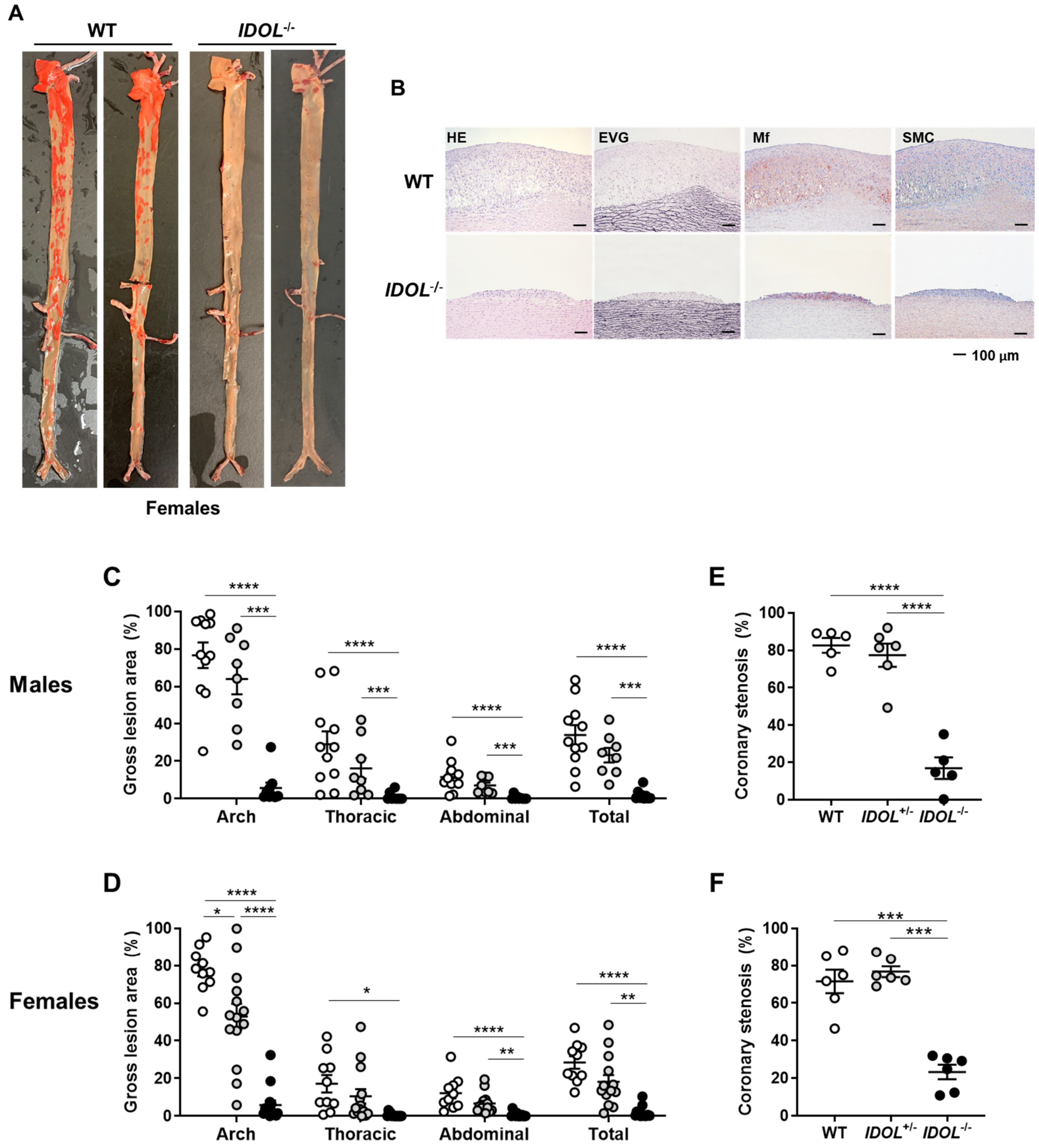


**Figure S4. Reduced atherosclerotic lesions in the IDOL KO rabbits is independent of sex.**

**A**, Representative pictures of aortic atherosclerosis in female WT and *IDOL*$^{-/-}$ rabbits after 16 weeks on HCD. Lipids in the arterial wall were stained en face with Sudan IV to evidence the atherosclerotic lesions. **B**, Representative pictures of the pathology of atherosclerotic lesions in the aortic arch from WT and *IDOL*$^{-/-}$ rabbits at endpoint. Samples were stained with hematoxylin and eosin (HE) and Elastica van Gieson (EVG). Macrophages (MF) and smooth muscle cells (SMC), were immunostained with RAM11 and HHF35 antibodies, respectively. Scale bar, 100μm. **C**, Atherosclerotic lesions in the aorta (as percentage) determined as in Figure 3B from males, compared across the three genotypes. WT, n=11; *IDOL*$^{+/-}$, n=8; *IDOL*$^{-/-}$, n=9. **D**, Atherosclerotic lesions in the aorta as in C, from females. WT, n=10; *IDOL*$^{+/-}$, n=14; *IDOL*$^{-/-}$, n=14. **E**, Dot plot of coronary atherosclerosis calculated as in Figure 3C in male rabbit. WT, n=5; *IDOL*$^{+/-}$, n=6; *IDOL*$^{-/-}$, n=5. **F**, Data as in E, from female rabbits. WT, n=6; *IDOL*$^{+/-}$, n=6; *IDOL*$^{-/-}$, n=6. Data are shown as mean±SEM. (C, F) 2-way ANOVA with Tukey's multiple comparison test. *p<0.05, **p<0.01, ***p<0.001, and ****p<0.0001.

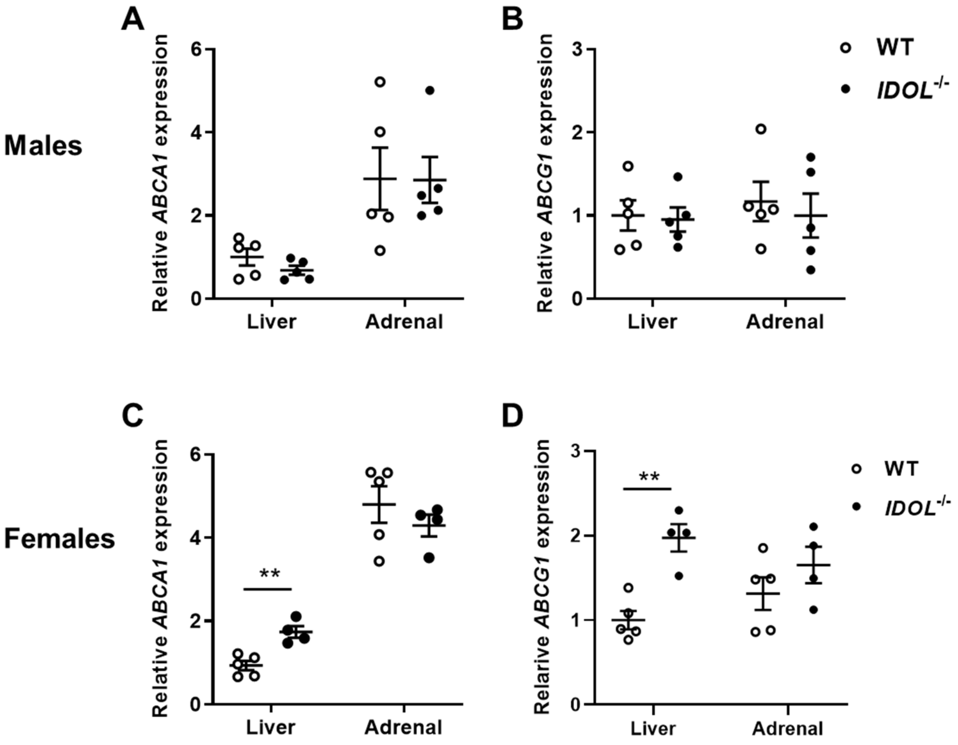


**Figure S5. Increased expression of cholesterol efflux genes in female liver.** Relative expression of ATP Binding Cassette Subfamily A Member 1 (*ABCA1*) and ATP Binding Cassette Subfamily G Member 1 (*ABCG1*), normalized by 18S, in the liver and adrenal gland from rabbits on HCD for 16 weeks. **A**, *ABCA1* and **B**, *ABCG1*, in males. **C**, *ABCA1* and **D**, *ABCG1*, in females. WT, n=10, (males, 5; females, 5); *IDOL*$^{-/-}$, n=9, (males, 5; females, 4). Data normalized to liver WT set as 1 and expressed as mean±SEM. Unpaired Students' t-test or Mann–Whitney U test, depending on the normality in distribution; each tissue compared independently by genotype. **p<0.01.

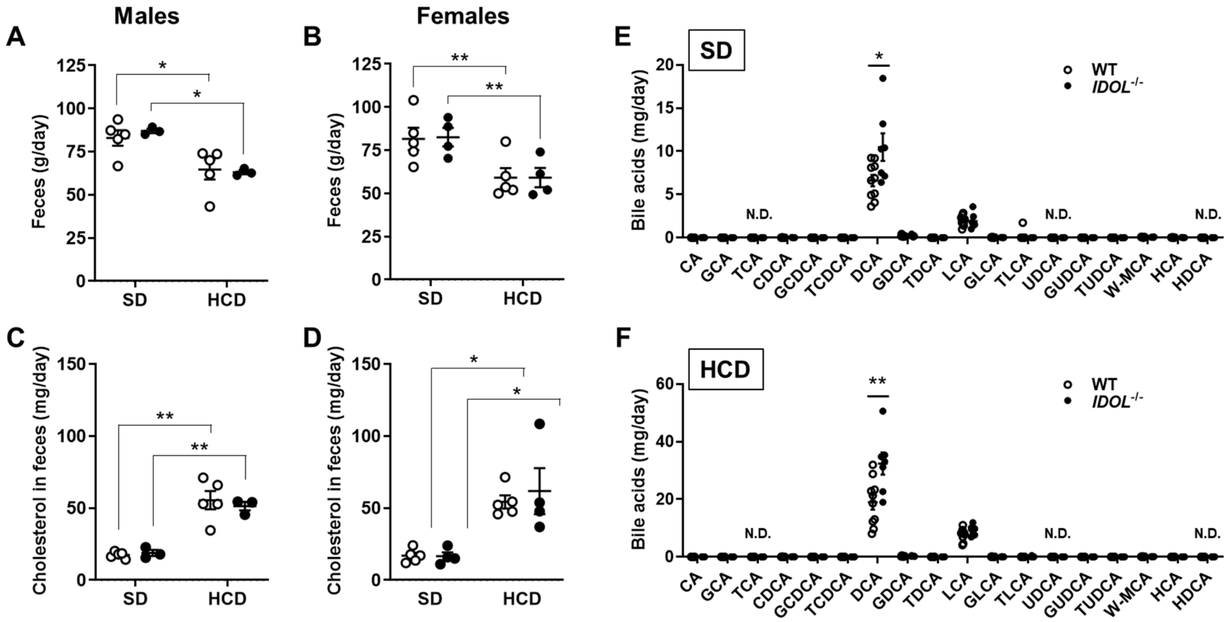


**Figure S6. Fecal cholesterol and bile acids excretion in the IDOL$^{-/-}$ rabbits.** Average daily weight of feces from **A**, males and **B**, females. Cholesterol in feces from **C**, males and **D**, females. Bile acids in feces from rabbits on **E**, SD and **F**, HCD (sexes combined) Deoxycholic acid (DCA), lithocholic acid (LCA). WT, n=10 (male, 5; female, 5); *IDOL*$^{-/-}$, n=7 (male, 3; female, 4). All data are shown as mean±SEM. (A , B, C, D) Two-way ANOVA with Sidak's multiple comparisons test. (E, F) Unpaired Students' t test for each bile acid comparing by genotype. * p<0.05, **p<0.01.
